# The anaphase-promoting complex regulates the degradation of the inner nuclear membrane protein Mps3 in budding yeast

**DOI:** 10.1101/415844

**Authors:** Bailey A. Koch, Hui Jin, Robert J. Tomko, Hong-Guo Yu

**Affiliations:** Department of Biological Science, the Florida State University, Tallahassee FL 32306; Department of Biomedical Sciences, the Florida State University, Tallahassee FL 32306

**Author notes:** Correspondence to: Hong-Guo Yu.

## Abstract

The nucleus is enclosed by the inner nuclear membrane (INM) and the outer nuclear membrane (ONM). While the ONM is continuous with the endoplasmic reticulum (ER), the INM is independent and separates the nucleoplasm from the ER lumen. To maintain INM homeostasis, proteins mislocalized to the INM are degraded by the ER-associated protein degradation (ERAD) pathway. However, the mechanism for turnover of resident INM proteins is less clear. Here we show that the anaphase-promoting complex/cyclosome (APC/C), an E3 ubiquitin ligase, regulates the degradation of Mps3, a conserved integral protein of the INM. Turnover of Mps3 requires the ubiquitin-conjugating enzymes Ubc7 and Ubc6, but not the three known ERAD ubiquitin ligases, Doa10, Hrd1, and the Asi1-Asi3 complex. Using a genetic approach, we have found that Cdh1, a coactivator of APC/C, modulates Mps3 stability. APC/C controls Mps3 degradation through Mps3’s N-terminus, which resides in the nucleoplasm and possesses two putative APC/C-dependent destruction motifs. Accumulation of Mps3 at the INM impairs nuclear morphological changes and cell division. Our findings therefore reveal an unexpected mechanism of APC/C-mediated protein quality control at the INM that coordinates nuclear morphogenesis and cell-cycle progression.

## Introduction

The nucleus is enclosed by two membranes that demarcate the nucleoplasm from the cytoplasm. The outer nuclear membrane (ONM) is continuous with the endoplasmic reticulum (ER), whereas the inner nuclear membrane (INM), which harbors hundreds of proteins (Ungricht and Kutay, 2015; Smoyer et al., 2016), interacts with the nucleoplasm. INM-localized proteins regulate a diverse range of nuclear activities that include chromosome movement, gene expression, and signal transduction. Nascent INM proteins are synthesized at the ER, transported through the nuclear pore complex, and then anchored at the INM (Katta et al., 2014; Ungricht and Kutay, 2015). Abnormal accumulation of INM proteins, such as the integral membrane protein SUN1, has been linked to the pathogenesis of progeric and dystrophic laminopathies in mammals (Chen et al., 2012; Burke and Stewart, 2014). But how homeostasis of resident INM proteins is achieved in order to maintain proper INM function remains to be further elucidated.

The ER-associated protein degradation (ERAD) pathway regulates the turnover of many ER proteins by marking them for proteasome degradation (Vembar and Brodsky, 2008; Zattas and Hochstrasser, 2015). ERAD acts in a step-wise manner which involves the target protein being polyubiquitylated by the joint actions of an E2 ubiquitin-conjugating enzyme and an E3 ubiquitin ligase. Two partially redundant E2 enzymes, Ubc6 and Ubc7, are known to function with one of the three independent ERAD E3 ligases, Doa10, Hrd1, and the Asi1-Asi3 complex in budding yeast, thus forming three separate branches of the ERAD pathway (Bordallo and Wolf, 1999; Swanson et al., 2001; Carvalho et al., 2006; Foresti et al., 2014; Khmelinskii et al., 2014). Because of the close association of the nuclear envelope to the ER, it is not surprising that ERAD also regulates protein quality control at the INM.

Recent work from budding yeast has shown that the Asi1-Asi3 protein complex in particular acts in concert with Ubc6 and Ubc7 to polyubiquitinate INM proteins that are sorted for proteasome degradation (Foresti et al., 2014; Khmelinskii et al., 2014), thereby defining an ERAD branch that operates specifically at the INM. However, the Asi proteins, including Asi1, Asi2 and Asi3, are not conserved in mammals, and genome-wide proteomic analysis in budding yeast has shown that they are mostly responsible for the degradation of mislocalized INM proteins (Foresti et al., 2014; Khmelinskii et al., 2014). Noticeably, being a resident INM protein itself, Asi1 is highly unstable and subject to proteasome degradation, but the responsible E3 ligase for Asi1 turnover remains unknown (Pantazopoulou et al., 2016). These observations indicate that additional E3 ligases function at the INM to regulate protein turnover.

We show here that the anaphase promoting complex/cyclosome (APC/C), an E3 ubiquitin ligase best known for its role in controlling cell-cycle progression (Irniger et al., 1995; King et al., 1995; Sudakin et al., 1995), regulates the degradation of the SUN-domain protein Mps3, an integral INM protein and an essential component of the linker of the nucleoskeleton to cytoskeleton (LINC) complex (Jaspersen et al., 2002; Conrad et al., 2007). Using a genetic approach, we show that APC/C mediates Mps3 degradation through Mps3’s N-terminus, which resides in the nucleoplasm and possesses two putative APC/C-dependent degradation motifs. Accumulation of Mps3 at the INM impairs nuclear morphological changes and cell division. Our work reveals that APC/C-mediated protein quality control at the INM coordinates nuclear morphogenesis with cell cycle progression.

## Results

### Degradation of Mps3 is regulated by the ubiquitin-proteasome system

We and others have shown previously that the INM-localized protein Mps3 regulates centrosome duplication and separation in budding yeast (Jaspersen et al., 2002; Friederichs et al., 2011; Li et al., 2017). Here we seek to determine how Mps3 is subject to protein degradation during the cell cycle. We generated an N-terminal V5-tagged allele, *P*_*MPS3*_*-V5-MPS3* (Fig 1A) which is under the control of its endogenous promoter and serves as the only copy of *MPS3* in the yeast genome. To determine the half-life of Mps3, we used cycloheximide (CHX), a potent inhibitor of protein biosynthesis, and performed cycloheximide-chase experiments (Fig 1A and see below). By western blotting, we found that V5-Mps3 exhibited a half-life of approximately 45 minutes when cells proliferated in rich medium (Fig 1B), demonstrating that Mps3 is an unstable protein.

**Figure 1.**
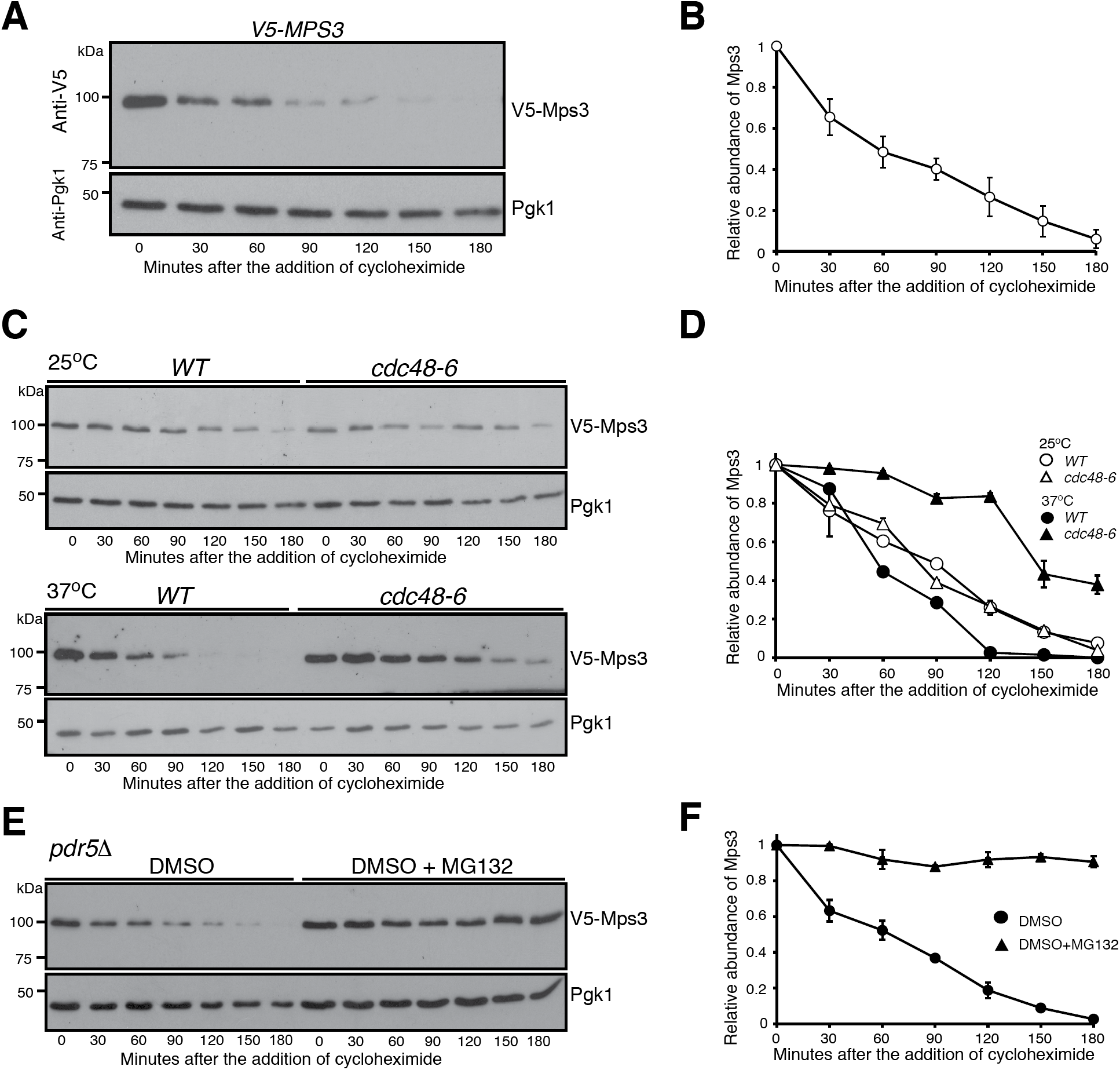
Proteasome-dependent degradation of Mps3. (**A** and **B**) Cycloheximide (CHX)-chase experiment showing Mps3 half-life. Yeast cells were grown to the exponential phase, then CHX was added to the culture media. Cell aliquots were withdrawn at indicated times, and protein extracts were prepared for western blotting. An anti-V5 antibody was used to probe V5-Mps3. The level of Pgk1 serves as a loading control. Time zero is the point of CHX addition. Quantification of Mps3 protein abundance is shown in panel B. Error bars represent the standard deviation from the mean of 7 biological replicates. (**C** and **D**) Cdc48 regulates Mps3 degradation. Cells were grown at 25°C to the exponential phase and CHX was then added as shown in A. To inactivate *cdc48-6*, cells were shifted to 37°C for 1 hour before the addition of CHX. Quantification of Mps3 protein abundance in shown in D. Samples were analyzed as in B; error bars represent the standard deviation from the mean of biological replicates (n=3). Notably, the half-life of Mps3 at the nonpermissive temperature increased more than 2 fold. (**E** and **F**) The proteasome is responsible for Mps3 turnover. Protein extracts and western blot were prepared as in A. The use of the *pdr5δ* allele allows yeast to uptake MG132, which is dissolved in DMSO. Quantification of Mps3 protein abundance in shown in F. Samples were analyzed as in B with error bars representing the standard deviation from the mean of biological replicates (n=2). Note that Mps3 is stabilized in the presence of MG132.

Because Mps3 is an integral membrane protein of the INM (Jaspersen et al., 2002), we determined whether Mps3 is subject to ERAD regulation. Although there are multiple ERAD branches, each utilizes the AAA-ATPase Cdc48, which facilitates the extraction of membrane proteins and is critical for transporting ubiquitinated substrates to the proteasome (Rabinovich et al., 2002; Schuberth and Buchberger, 2005). We therefore determined whether Mps3 is regulated by Cdc48 (Fig 1C). We used a temperature-sensitive allele, *cdc48-6,* to inactivate Cdc48 at 37°C. At the permissive temperature (25°C), the half-life of Mps3 was comparable in wild-type and *cdc48-6* cells (Fig 1C and 1D). In contrast, the Mps3 protein level stabilized in *cdc48-6* cells at the restrictive temperature (37°C), with a half-life about 150 minutes (Fig 1D), a 3-fold increase. This finding suggests that Mps3 is likely subject to ERAD regulation. To further confirm that Mps3 is degraded by the proteasome, we used the cell-permeable drug MG132 to inhibit proteasome activity. In order for yeast cells to retain MG132 and thereby keep the proteasome at an inactive state, the plasma membrane ABC transporter, encoded by *PDR5*, was removed (Leppert et al., 1990). Noticeably, in *pdr5Δ* cells treated with MG132, Mps3 became highly stable, showing a half-life of more than 180 minutes (Fig 1E and 1F). We therefore conclude that the ubiquitin proteasome system regulates Mps3 degradation.

### Mps3 is degraded by nucleus-localized proteasomes

The N-terminal domain of Mps3 is positioned in the nucleoplasm (Jaspersen et al., 2002; Li et al., 2017); we therefore asked whether Mps3 is degraded by nucleus-localized proteasomes. To deplete proteasomes from the nucleus, we used the anchor-away system to force proteasomes to relocate to either the cytoplasm or the plasma membrane (Haruki et al., 2008; Nemec et al., 2017 and Fig 2A). Of note, tethering the proteasome to the anchor protein, which is mediated by rapamycin-triggered dimerization of FRB and FKBP, is conditional (Haruki et al., 2008) (Fig 2A), and the *TOR1-1* mutation renders yeast cells resistant to rapamycin (Helliwell et al., 1994). We found that the half-life of Mps3 was not altered upon the addition of rapamycin, which served as a control in our experiments (Fig 2B and C). However, upon the addition of rapamycin and by way of Rpn11-FRB and Rpl13-FKBP interaction, proteasomes were sequestered to the cytoplasm (Fig 2D). As a result, the half-life of Mps3 reached 150 minutes, a 3-fold increase over the wild type (Fig 2B and C), demonstrating that degradation of Mps3 was impaired when proteasomes were depleted from the nucleus. Similarly, anchoring proteasomes to the plasma membrane also increased the half-life of Mps3 to above 150 minutes (Fig 2E, F and G). Taken together, our findings suggest that the bulk of Mps3 is degraded by nuclear proteasomes.

**Figure 2.**
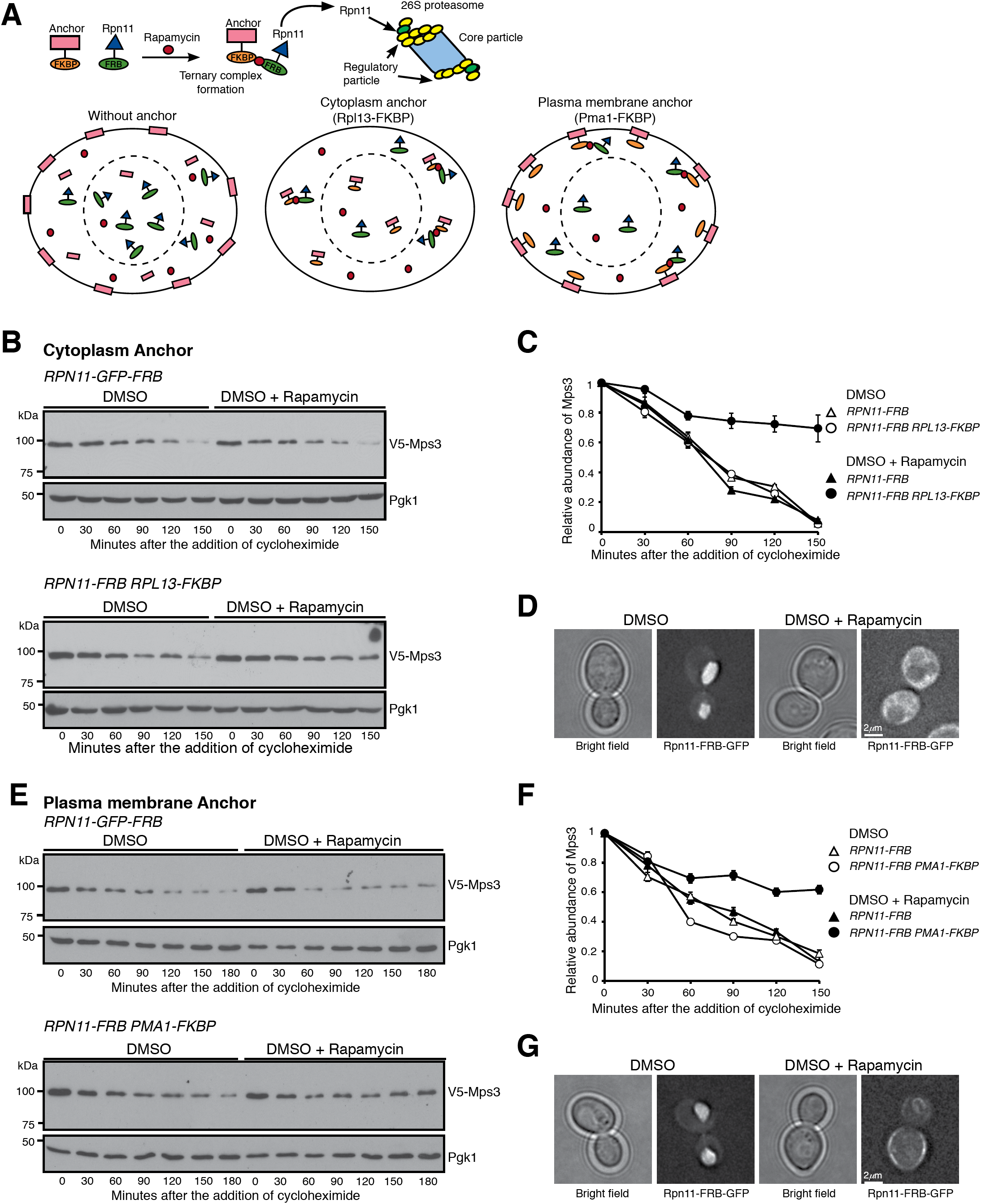
Nucleus-localized proteasome regulates Mps3 degradation. **(A)** A schematic representation of forced proteasome localization using the anchor-away system. Briefly, Rpn11, a subunit of the 26S proteasome was tagged with FRB, the presence of rapamycin induces a ternary complex formation between Rpn11-FRB and the FKBP-tagged anchor, either at the cytoplasm (Rpl13-FKBP) or at the plasma membrane (Pma1-FKBP). **(B)** Forced localization of proteasomes to the cytoplasm. Yeast cells were prepared for CHX chase, and rapamycin dissolved in DMSO was added to relocate the proteasome to the cytoplasm. **(C)** Quantification of Mps3 degradation as shown in B. Error bars represent the standard deviation from the mean of biological replicates (n=2). **(D)** Representative images showing forced localization of the proteasome to the cytoplasm. Note that Rpn11-GFP-FRB, became dispersed in the cytoplasm in the presence of rapamycin. **(E)** Forced localization of the proteasome to the plasma membrane. Samples were analyzed as in B. **(F)** Quantification of Mps3 degradation as shown in E. Error bars represent the standard deviation from the mean of biological replicates (n=2). **(G)** Representative images showing forced localization of the proteasome to the plasma membrane. Note that Rpn11-GFP-FRB was enriched at the cell periphery in the presence of rapamycin.

### The ERAD E2 enzymes Ubc7 and Ubc6 regulate Mps3 degradation

We sought to identify the E2 ubiquitin conjugating enzyme and the E3 ubiquitin ligase that are responsible for Mps3 degradation. We have reported previously that overproduction of Mps3 by way of *P*_*G*__*AL*_*-MPS3* leads to a slight growth defect in vegetative yeast cells (Li et al., 2017). Because Mps3 is an essential component of the INM and the SPB, we reasoned that impairing the degradation system specific to Mps3 would render a lethal phenotype when Mps3 is overproduced. From the yeast deletion collection library (ATCC-GSA-5), we pooled together deletions of nonessential yeast genes that encode either an E2 or an E3 or their interacting factors, and screened their genetic interactions with *P*_*GAL*_*-MPS3*, which overproduced Mps3 in the galactose medium (Fig 3A as an example). Of note, in the dextrose medium, the *GAL* promoter is repressed. We found that deletion of *UBC7*, which encodes an E2 for ERAD (Gilon et al., 1998), showed synthetic lethality with *P*_*GAL*_*-MPS3* when yeast cells were cultured in the galactose medium (Fig 3A and B). In addition, *DOA4*, which encodes the ubiquitin hydrolase that is responsible for recycling ubiquitin from proteasome-bound intermediates (Papa and Hochstrasser, 1993), also genetically interacted with *P*_*GAL*_*-MPS3* (Fig 3B). Importantly, the synthetic lethality observed in *P*_*GAL*_*-MPS3 doa4Δ* cells was suppressed upon the introduction of a high-copy-number plasmid that expressed ubiquitin (Fig 3B). These findings indicate that Ubc7 is a putative E2 enzyme for Mps3 degradation, and support the idea that degradation of Mps3 is mediated by the ERAD ubiquitin-proteasome system.

**Figure 3.**
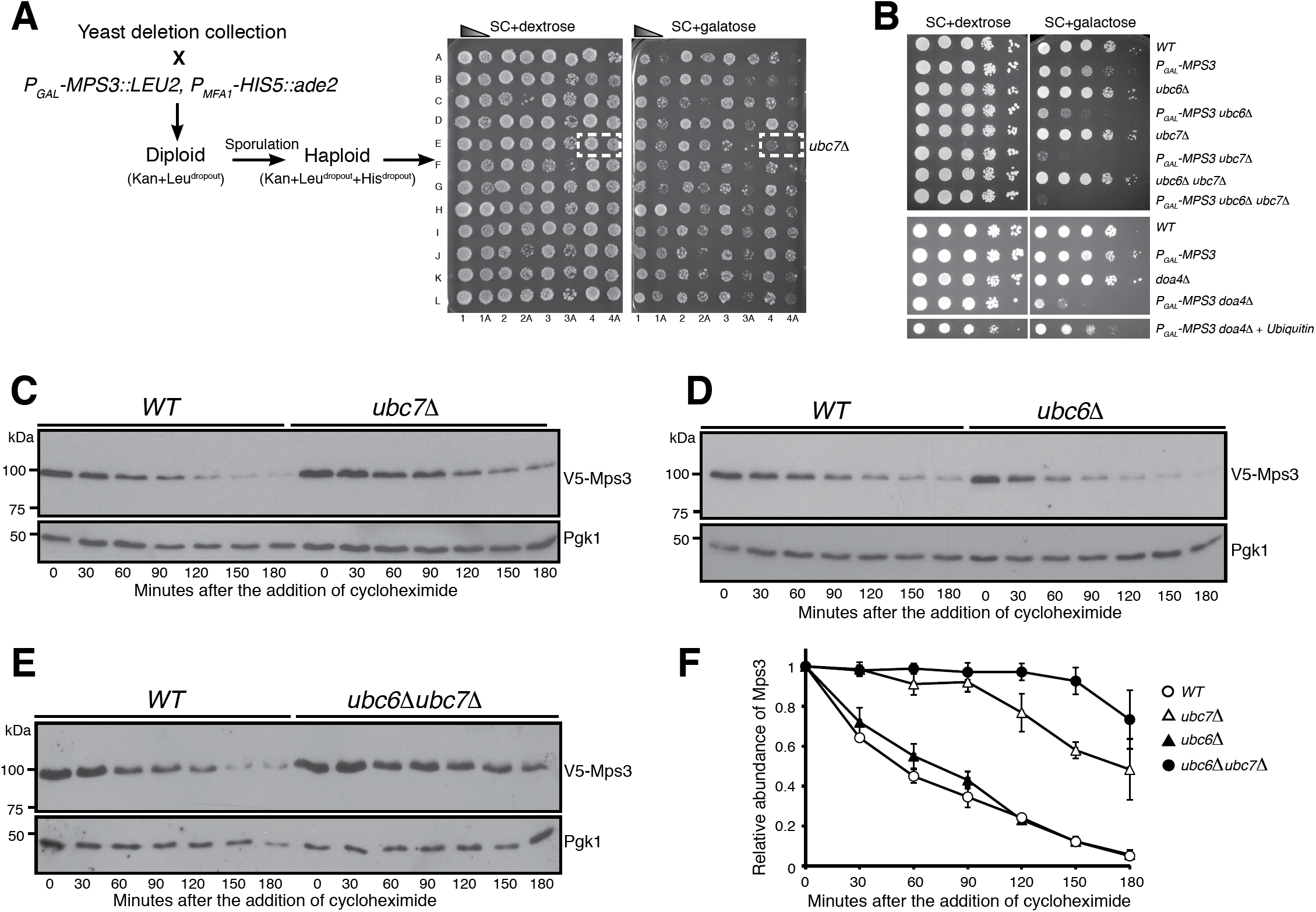
Ubc7 is a key regulator of Mps3 degradation. **(A)** A representation of the genetic screen performed utilizing the yeast deletion collection. Briefly, a deletion collection specific for genes encoding known ubiquitin conjugating enzymes, ubiquitin ligases, and their regulators was crossed to a strain containing an *MPS3* overexpression plasmid under the galactose inducible promoter. Selective media allowed for the selection of haploid yeast cells containing both *P*_*GAL*_*-MPS3* and a gene deletion. This method enabled the identification of putative genetic interactors of *MPS3*. The synthetic lethality observed with *ubc7δ, doa4δ*, and others was further tested to confirm the genetic interaction. (**B**) *MPS3* genetically interacts with *UBC7* and *DOA4*. To determine cell viability, yeast cells were grown overnight in YPD liquid medium to reach saturation, 10 fold diluted, spotted onto SC plates with either 2% dextrose or 2% galactose, and then incubated at 30°C for about two days. Overexpression of *MPS3* yielded a sick phenotype. The overexpression of *MPS3* coupled with *ubc6δ* cells showed a stronger sick phenotype while the *ubc7δ* and *ubc6δubc7δ* had a lethal phenotype. Notably, the similar near lethal phenotype observed when coupled with a *doa4δ* can be suppressed by the overexpression of ubiquitin. (**C**, **D** and **E**) Ubc7 regulates degradation of Mps3. WT, *ubc6δ, ubc7δ*, and *ubc6δubc7δ* yeast cells were prepared for CHX chase as in Figure 1A. **(F)** Quantification of the relative abundance of Mps3 as shown in C, D, and E. Error bars represent the standard deviation from the mean of biological replicates (n=2).

To determine whether Ubc7 governs Mps3 protein stability, we determined the half-life of Mps3 in *ubc7Δ* cells (Fig 3C). Indeed, in the absence of Ubc7, the half-life of Mps3 increased to more than 150 minutes (Fig 3C and see below). Another E2 enzyme involved in ERAD, Ubc6, is partially redundant to Ubc7 (Chen et al., 1993). We therefore determined Mps3’s half-life in *ubc6Δ* cells (Fig 3D). Deletion of *UBC6* alone did not change Mps3’s stability when compared to wild type (Fig 3D). However, removal of both Ubc6 and Ubc7 (*ubc6Δ ubc7Δ*) drastically increased Mps3’s half-life as compared to the *ubc7Δ* single mutant (Fig 3E and F). This finding indicates that in the absence of Ubc7, Ubc6 can partially serve as an alternative E2 for Mps3 degradation. We therefore conclude that the ERAD-specific E2 enzymes Ubc7 and, to a lesser degree, Ubc6 regulate Mps3 protein turnover.

### Mps3 is not regulated by the canonical ERAD-L/M/C E3 ubiquitin ligases

Both Ubc6 and Ubc7 are involved in ERAD-mediated protein quality control; we therefore hypothesized that the known ERAD E3 ubiquitin ligases, Hrd1, Doa10, and the Asi1-Asi3 complex (Bordallo and Wolf, 1999; Swanson et al., 2001; Carvalho et al., 2006; Foresti et al., 2014; Khmelinskii et al., 2014), are required for Mps3 degradation. To test our hypothesis, we used gene deletions of *HRD1, DOA10*, and *ASI3* to abolish their corresponding E3 enzymatic activity (Fig 4). Using cycloheximide chase, we observed that the degradation of Mps3 is independent of Hrd1 (Fig 4A and 4B), an E3 ubiquitin ligase responsible for retrotranslocating substrates from the ER to cytosol for degradation (Bordallo and Wolf, 1999; Baldridge and Rapoport, 2016). This result supports our finding that turnover of Mps3 takes place inside the yeast nucleus (Fig 2). The other two E3 enzymes, Doa10 and the Asi1-Asi3 complex, both have been implicated in degradation of certain INM-localized proteins (Swanson et al., 2001; Foresti et al., 2014; Khmelinskii et al., 2014). However, the half-life of Mps3 remained unchanged in *doa10Δ* or *asi3Δ*, or *doa10Δasi3Δ* double mutant cells (Fig 4C and D, data not shown). More importantly, removal of all the three known ERAD E3 enzyme activities had no noticeable impact on the half-life of Mps3 (Fig 4E and F). By contrast, a previously identified INM protein and Asi1-Asi3-complex substrate, Erg11 (Foresti et al., 2014), was stabilized when the activities of the three canonical ERAD E3 ligases were absent (Fig 4E and F). Taken together, our findings demonstrate that degradation of Mps3 is not governed by the three E3 ubiquitin ligases known for ERAD, and indicate that a different E3 ligase is responsible for regulating Mps3 protein stability at the INM.

**Figure 4.**
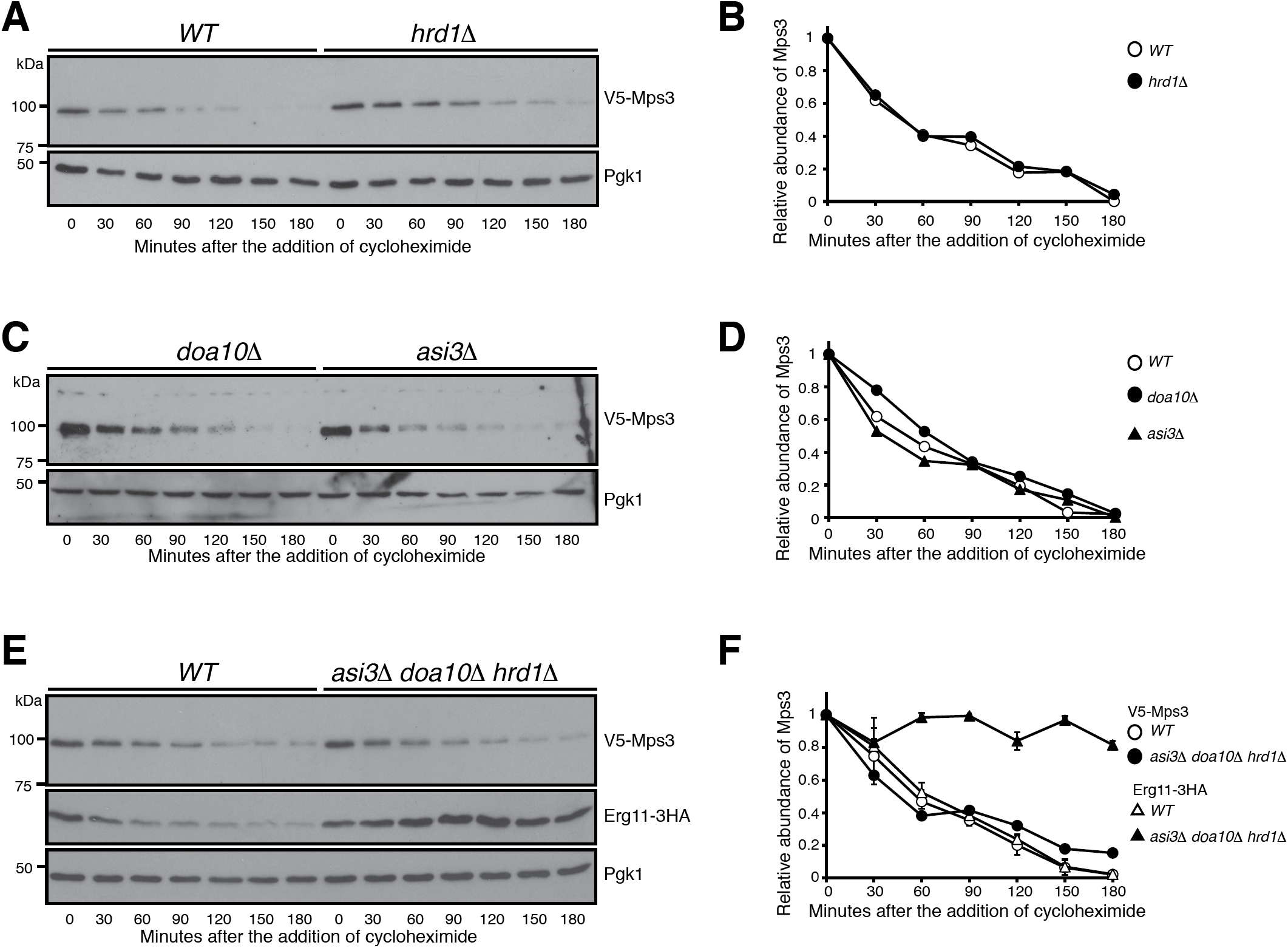
Mps3 degradation is independent of the canonical ERAD E3 ligases. **(A** and **B)** Canonical E3 ligase Hrd1 does not regulate Mps3 degradation. *WT* and *hrd1δ* yeast cells were prepared for CHX chase as in Figure 1A. Quantification of Mps3 protein abundance is shown in panel B. **(C** and **D)** Canonical E3 ligases Doa10 and Asi3 do not regulate Mps3 degradation. Yeast cells were prepared for CHX chase in a *doa10δ* or *asi3δ* background and were analyzed as in Figure 1A. Quantification of Mps3 protein abundance as shown in D. **(E** and **F)** The canonical E3 ligases involved in ERAD do not regulate Mps3 degradation through redundancy. *WT* and *asi3δdoa10δhrd1δ* yeast cells were prepared for CHX chase and analyzed as in Figure 1A. Erg11-3HA was used as a positive control and Pgk1 was probed for a loading control. Quantification of Mps3 relative protein abundance is shown in F. Error bars represent the standard deviation from the mean of biological replicates (n=2). Note there is no change in Mps3 half-life between the *WT* and *asi3δdoa10δhrd1δ* cells.

### The anaphase-promoting complex/cyclosome regulates Mps3 degradation

To search for the E3 ubiquitin ligase that is responsible for Mps3 degradation, we took a genetic approach similar to that described in Fig 3A to identify Mps3 interacting factors that regulate the activities of yeast E3 ligases. One of them, *CDH1*, which encodes an activator of the anaphase-promoting complex/cyclosome (APC/C) (King et al., 1995; Visintin et al., 1997; Zachariae et al., 1998), genetically interacted with *P*_*GAL*_*-MPS3*; deletion of *CDH1* caused a lethal phenotype when *MPS3* was overexpressed (Fig 5A). Cdh1 acts during late mitosis and at the G1 phase (Visintin et al., 1997); we therefore enriched yeast cells in G1 with alpha factor and performed cycloheximide-chase experiments to determine Mps3’s half-life (Fig 5B). Consistent with their genetic interaction, the half-life of Mps3 increased to more than 3 hours in the absence of Cdh1 (Fig 5B and C). Cdh1 has a paralog called Cdc20, which activates APC/C at the metaphase to anaphase transition during mitosis (Visintin et al., 1997). Because *CDC20* is an essential gene, we used a temperature-sensitive *cdc20-1* allele to inactivate Cdc20 at 37°C (Fig 5C and D). The half-life of Mps3 remained unchanged in *cdc20-1* cells at the restrictive temperature as in wild-type cells (Fig 5C and D). We therefore conclude that Cdh1, but not Cdc20, specifically regulates Mps3 turnover.

**Figure 5.**
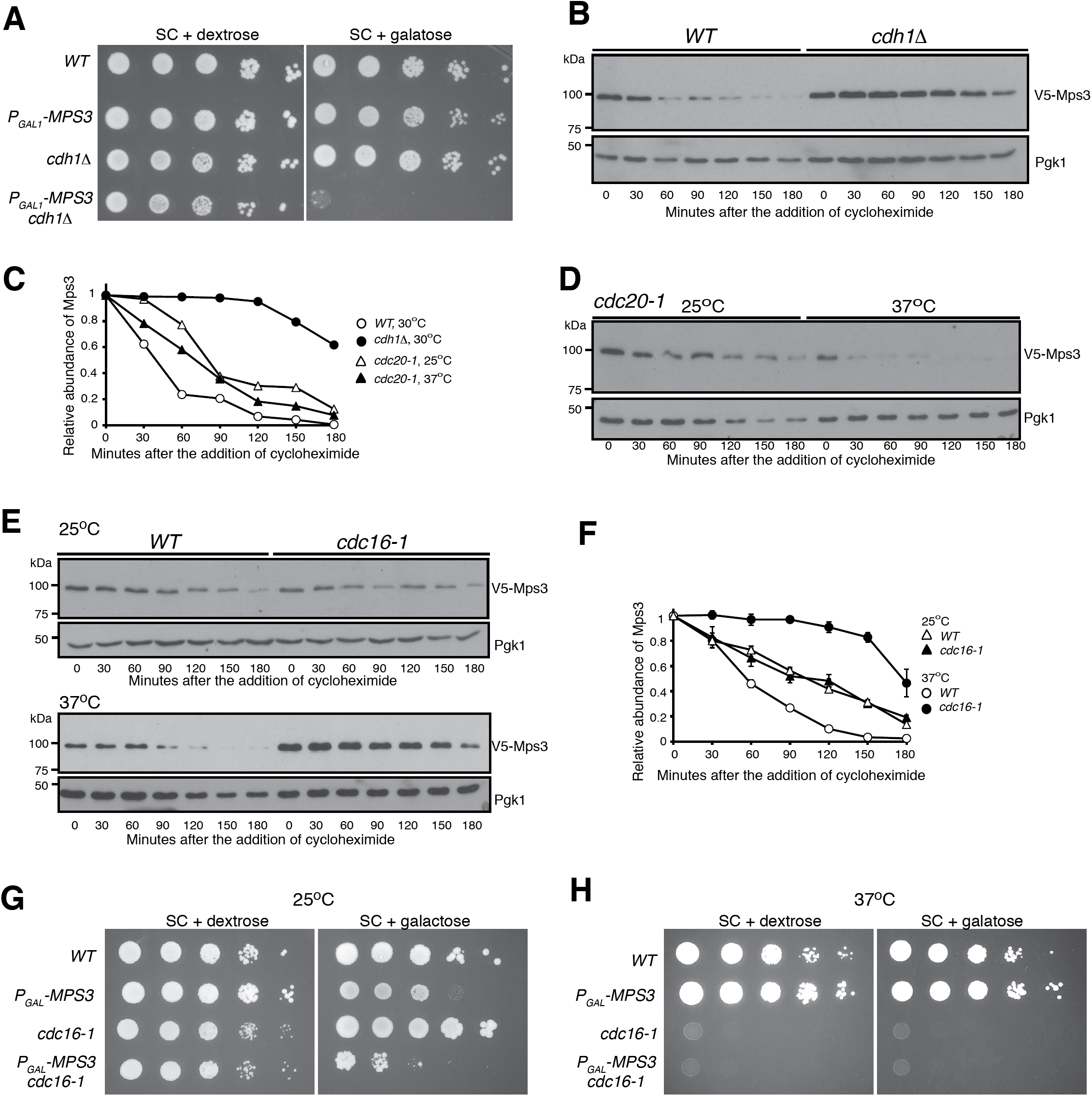
APC/C^Cdh1^-dependent regulation of Mps3 degradation. (**A**) *MPS3* genetically interacts with *CDH1*. To determine cell viability, yeast cells were grown overnight in YPD liquid medium to reach saturation, 10 fold diluted, spotted onto SC plates with either 2% dextrose or 2% galactose, and then incubated at 30°C for about two days. Overexpression of *MPS3* is lethal in *cdh1δ* cells. **(B)** The APC/C activator, Cdh1, regulates Mps3 degradation. The relative protein abundance of Mps3 in *WT* and *cdh1δ* cells was analyzed after arresting cells in G1 using alpha factor (10 μg/mL) for 2 hours and then CHX chase was performed as described in Figure 1A. (**C**) Quantification of the relative abundance of Mps3 as shown in B and D. Representative blots from two biological replicates of each experiment are shown. **(D)** Mps3 turnover is independent of Cdc20. Cells were grown at 25°C to the exponential phase and CHX was then added as shown in Figure 1A. To inactivate *cdc20-1*, cells were shifted to 37°C for 1-hour before the addition of CHX. **(E** and **F)** The APC/C subunit Cdc16 regulates Mps3 degradation. Cells were grown at 25°C to the exponential phase and CHX was then added as shown in Figure 1A. To inactivate *cdc16-1*, cells were shifted to 37°C for 1 hour before the addition of CHX. Quantification of Mps3 protein abundance is shown in F. Error bars represent the standard deviation from the mean of biological replicates (n=2). Notably, inactivation of the APC through the *cdc16-1* allele at the nonpermissive temperature resulted in a 3-fold increase in Mps3 half-life. (**G** and **H**) *MPS3* genetically interacts with *cdc16-1*. Overexpression of *MPS3* in *cdc16-1* cells showed a sick phenotype at the permissive temperature. Yeast cells were grown overnight in YPD liquid medium to reach saturation, 10-fold diluted, spotted onto SC plates with either 2% dextrose or 2% galactose, and then incubated at 25°C and 37°C for about two days.

We hypothesized that APC/C serves as the E3 ligase for Mps3 degradation. To test this hypothesis, we inactivated the APC/C using the temperature sensitive *cdc16-1* allele, the wild type of which encodes an essential subunit of the APC/C (Zachariae et al., 1996). By cycloheximide chase and western blotting, we found that Mps3 was stabilized when *cdc16-1* was inactivated at 37°C, reaching a half-life of 160 minutes, about a 4-fold increase over that of wild type (Fig 5E and F). As a control, we found that Asi1, another INM-localized protein whose responding E3 is currently unknown, was degraded at a similar kinetics in *cdc16-1* cells as compared to wild-type cells (Supplemental Fig 1), demonstrating the specificity of Cdc16 for Mps3 degradation. On the basis of this observation, we hypothesized that a genetic interaction would exist between *MPS3* and *CDC16*. To test this idea, we overproduced Mps3 in *cdc16-1* cells (Fig 5G and H). Even at the permissive temperature (25°C), we observed a severe growth defect when *P*_*GAL*_*-MPS3 cdc16-1* cells were cultured in the galactose medium (Fig 5G). Taken together, our findings indicate that APC/C^Cdh1^ regulates Mps3 degradation.

### The N-terminus of Mps3 possesses two putative APC/C-dependent destruction motifs

To mediate protein degradation, APC/C recognizes two well-defined destruction motifs, the D and KEN boxes, on its substrates (Glotzer et al., 1991; Pfleger and Kirschner, 2000). Deletion of amino acids from 64 to 93 (*mps3-NC*) drastically increased Mps3 protein stability in yeast meiosis (Li et al., 2017). This region harbors a KEN sequence from amino acids 67 to 73, whereas the amino acids from 76 to 84 loosely fit the consensus sequence of the D box (Fig 6A). To determine whether the putative KEN or D box, or both regulate Mps3 protein stability, we generated individual and double mutants: *mps3-3A,* in which the KEN sequence was replaced by three alanines; *mps3-2D*, in which two lysine residues at positions 76 and 76 were replaced by two glutamates; and *mps3-3A2D*, a double mutation (see diagram in Fig 6A). Overexpression of *mps3-3A2D*, but not *mps3-3A* or *mps3-2D*, rendered a lethal phenotype (Fig 6A and Supplemental Fig 2). Similarly, in either *mps3-3A* or *mps3-2D* mutant cells, whose expressions were under the control of the endogenous promoter, the half-life of Mps3 remained at or close to the wild-type level (Supplemental Fig 2). In contrast, in the double mutant *mps3-3A2D*, the half-life of Mps3 increased to more than 3 hours (Fig 6C and D). In addition, we found that the half-life of Mps3 dramatically increased in *mps3-NC* cells, in which the two putative destruction motifs both are absent (Fig 6E and F). These findings indicate that Mps3 possesses two putative destruction motifs, and the KEN and D boxes are somehow redundant in regulating Mps3 degradation.

**Figure 6.**
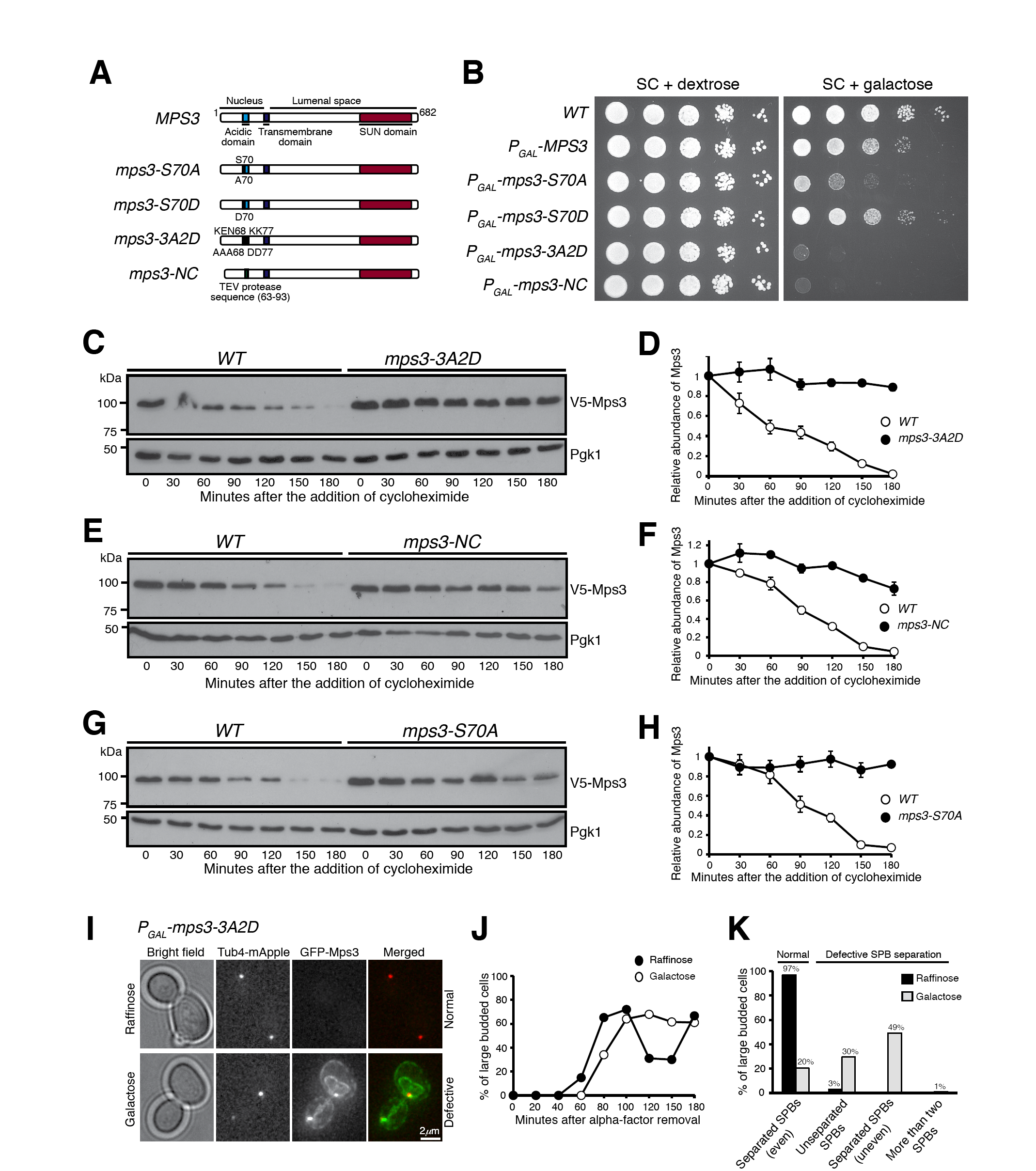
The N-terminus of Mps3 possesses two putative destruction motifs. **(A)** A schematic representation of Mps3 protein domains and subsequent mutations introduced. **(B)** The N-terminus of Mps3 is crucial for cell viability. Yeast cells were grown overnight in YPD liquid medium to reach saturation, 10 fold diluted, spotted onto SC plates with either 2% dextrose or 2% galactose, and then incubated at 30°C for about two days. Overexpression of a phosphorylation mutant (S70A) produces a very sick phenotype that can be repressed by the phosphomimetic (S70D) version. Amino acids 63-93 appear to be a key region of Mps3, where removal and specific point mutations results in a complete synthetic lethality. (**C-H**) The N-terminus of Mps3 regulates protein stability. WT, *mps3-3A2D* (**C** and **D**), *mps3-NC* (**E** and **F**), and *mps3-S70A* (**G** and **H**) yeast cells were prepared for CHX chase and analyzed as described in Figure 1A. Quantification of Mps3 protein abundance is shown in D, F, and H. Error bars represent the standard deviation from the mean of biological replicates (n=2). Note that each of these mutations results in a severe impairment of Mps3 degradation, with observed half-lives well over 180 minutes. (**I-K**) Accumulation of *P*_*GAL1*_*-mps3-3A2D* results in failed SPB separation or misseparation. Yeast cells were grown in raffinose medium and arrested at G1 with alpha-factor. Galactose was added 30 minutes prior to removal of alpha-factor to induce the *GAL1* promoter. Aliquots were withdrawn and prepared for live-cell fluorescence microscopy.Tub4-RFP marks the spindle pole body (SPB). Representative images of *mps3-3A2D* cells grew in raffinose or galactose are shown in I. Budding index is shown in J. Cell aliquots were withdrawn at indicated times and budding morphology was determined by phase contrast microscopy. Quantification of SPB separation is shown in K. Four categories of SPB separation in large budded cells were classified, the first being normal SPB separation (even), and the remaining three being types of defective SPB separation: unseparated SPBs, uneven separation of SPBs, and more than two SPBs. Tub4-RFP marks the SPB.

We have reported previously that Mps3 is phosphorylated at S70 (Li et al., 2017). Because protein phosphorylation regulates ubiquitination (Hunter, 2007), we addressed whether S70 phosphorylation regulates Mps3 stability using the phosphomutant *MPS3-S70A* (Li et al., 2017). Overproduction of Mps3-S70A, but not Mps3-S70D, which is a phosphomimetic, caused a severe growth defect (Fig 6B and Li et al., 2017). In the absence of S70 phosphorylation, Mps3 became highly stable, with a half-life of more than 3 hours (Fig 6G and H). Taken together, the above findings provide further evidence that posttranslational modification regulates Mps3 degradation.

To determine the biological significance of Mps3 accumulation at the INM, we used alpha factor to synchronize yeast cells at G1 and determined nuclear morphology and cell-cycle progression in cells with excessive Mps3-3A2D (Fig 6I-K). Upon G1 release by removal of the alpha factor, we overexpressed *P*_*GAL*_*-MPS3-3A2D* in the galactose medium (Fig 6I and Supplemental Fig 3). Overproduced Mps3-3A2D localized to the SPB and accumulated at the nuclear periphery (Fig 6I and supplemental Fig 3), forming extended membrane structures, a phenotype similar to that we reported previously of the *mps3-NC* allele (Li et al., 2017). On the basis of budding index and the dynamics of Tub4-marked SPBs, we determined that mutant cells overproducing Mps3-3A2D showed delayed cell-cycle progression (Fig 6J and K). In large budded cells at mitosis, 80% of them displayed SPB separation defects (Fig 6K). These findings explain the cell lethality caused by Mps3-3A2D overproduction.

### The N-terminus of Mps3 is both necessary and sufficient for APC/C^Cdh1^-mediated protein degradation

To determine if the N-terminus of Mps3 is sufficient for APC/C-mediated protein turnover, we engineered a Mps3(1-94)-Heh2 hybrid protein by grafting the N-terminus of Mps3 to Heh2 (Fig 7A), another INM-localized protein (King et al., 2006). We used the galactose medium to induce the expression of *P*_*GAL*_*-HEH2* and *P*_*GAL*_*-MPS3(1-94)-HEH2,* and determined protein stability by cycloheximide chase (Fig 7B-I). Heh2 was a relatively stable protein with a half-life of more than 180 minutes, whereas Mps3(1-94)-Heh2 displayed a half-life of less than 30 minutes (Fig 7B-E), demonstrating that the N-terminus of Mps3 is sufficient to mediate the degradation of the hybrid protein. Crucially, in the absence of Cdh1, Mps3(1-94)-Heh2 restored protein stability to a level similar to that of Heh2 (Fig 7D and E). As a control, overproduced Mps3 by way of *P*_*GAL*_*-GFP-MPS3* responded similarly to Cdh1 regulation just as the endogenous Mps3 (Fig 7F and see above). To determine whether APC/C regulates the degradation of the fusion protein, we used the *cdc16-1* allele to inactivate APC/C as shown in Fig 5E. At the restrictive temperature and in the absence of APC/C activity, Mps3(1-94)-Heh2 became highly stable (Fig 7H and I). These findings support our conclusion that the N-terminus of Mps3 specifically responds to APC/C^Cdh1^-mediated protein degradation.

**Figure 7.**
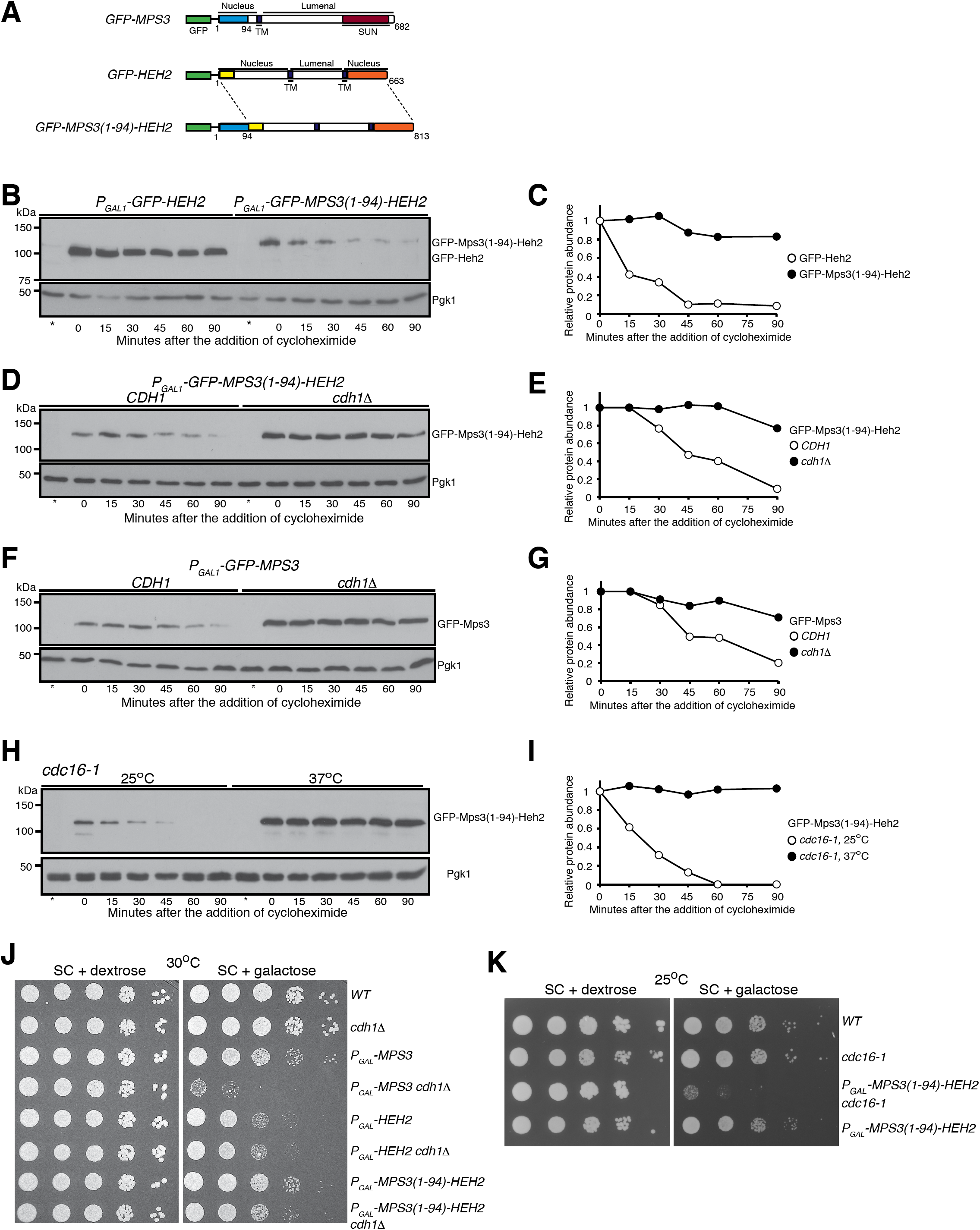
The N-terminus of Mps3 is necessary and sufficient for APC/C^Cdh1^-dependent protein degradation. **(A)** A schematic representation of GFP-Mps3, GFP-Heh2, and the fusion protein GFP-Mps3(1-94)-Heh2 constructs. (**B** and **C**) The N-terminus of Mps3 is sufficient for triggering the degradation of Mps3(1-94)-Heh2. CHX chase is as described in Figure 1A. To induce protein production, yeast cells were grown in galactose medium for ∼3 hours before the addition of CHX. Quantification of protein abundance is shown in panel C. Note that the fusion protein Mps3(1-94)-Heh2 has a half-life of about 30 minutes. (**D** and **E**) Cdh1 regulates Mps3(1-94)-Heh2 degradation. CHX-chase experiments were performed as in **B**. Quantification of protein abundance is shown in E. (**F** and **G**) Cdh1 regulates the degradation of overproduced Mps3. CHX-chase experiments were performed as in B. Quantification of protein abundance is shown in G. (**H** and **I**) Cdc16 regulates the degradation of the fusion protein Mps3(1-94)-Heh2. CHX-chase experiments were performed as in Figure 5E. To inactivate *cdc16-1*, cells were shifted to 37°C for 1-hour before the addition of CHX. Quantification of Mps3 protein abundance is shown in I. (**J** and **K**) Genetic evidence showing the N-terminus of Mps3 regulates protein stability. Yeast cells were grown overnight in YPD liquid medium to reach saturation, 10 fold diluted, spotted onto SC plates with either 2% dextrose or 2% galactose, and then incubated at 30°C (**J**) and 25°C (**K**) for about two days.

**Figure 8.**
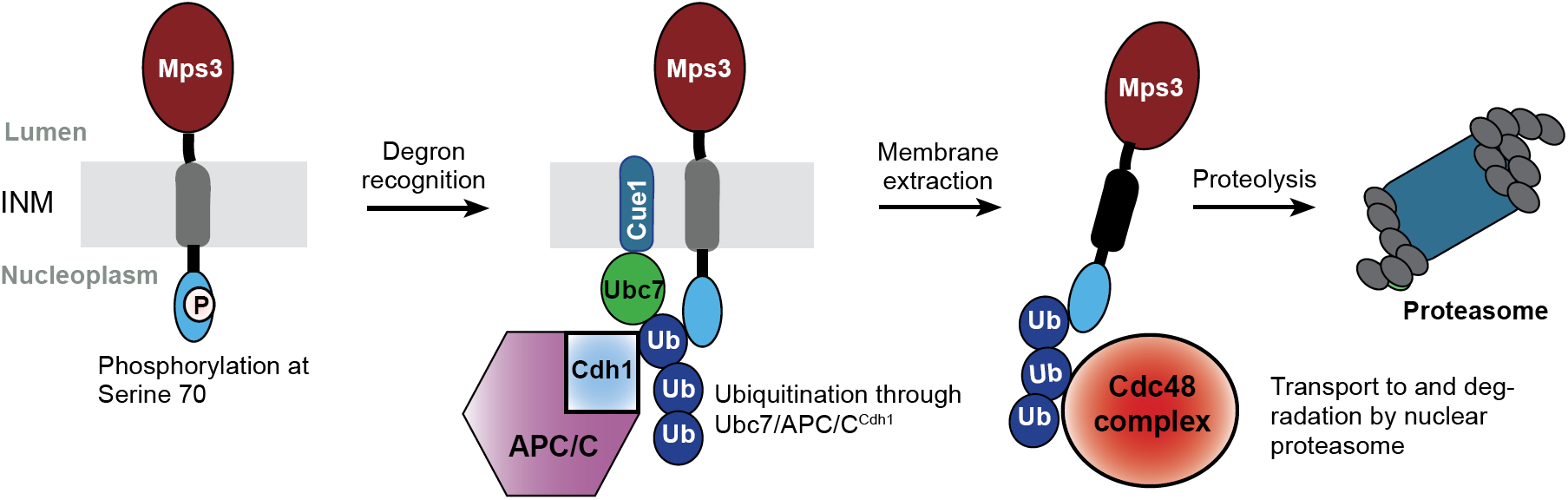
Model for APC/C-dependent Mps3 degradation. Mps3 is oligomerized at the INM (Inner nuclear membrane); for simplicity, only one copy of Mps3 is shown. Ub, ubiquitin.

Finally, overproduction of Heh2 by means of *P*_*GAL*_*-HEH2* caused a slow growth defect in yeast cells plated on the galactose medium (Fig 7J). In contrast, *P*_*GAL*_*-MPS3(1-94)-HEH2* cells displayed close to normal growth (Fig 7J), supporting the idea that the Mps3 N-terminus promotes the degradation of the fusion protein Mps3(1-94)-Heh2. In the absence of Cdh1, the fusion protein was stabilized (Fig 7C), and *P*_*GAL*_*-MPS3(1-94)-HEH2* cells grew slower in the galactose medium (Fig 7J). Crucially, even at the permissive temperature, *P*_*GAL*_*-MPS3(1-94)-HEH2 cdc16-1* cells failed to grow in the galactose medium (Fig 7K). Taken together, our findings demonstrate that the N-terminus of Mps3 is both necessary and sufficient for targeted protein degradation mediated by APC/C^Cdh1^.

## Discussion

In this report, we have shown that the degradation of the resident INM protein Mps3 is regulated by the E3 ubiquitin ligase APC/C in concert with the E2 ubiquitin-conjugating enzymes Ubc7 and Ubc6, thereby defining a new branch of the ERAD pathway for protein quality control at the nuclear envelope. In contrast to the three known ERAD E3 ligases, Doa10, Hrd1, and the Asi1-3 complex, all of which contain integral membrane proteins (Vembar and Brodsky, 2008; Zattas and Hochstrasser, 2015), APC/C does not possess membrane components and is best known for its role in degrading soluble proteins and regulating cell cycle progression (Pines, 2011; Davey and Morgan, 2016). Our work therefore extends the role of APC/C in modulating the turnover of an integral membrane protein. It also enriches the repertoire of the ERAD pathway for protein quality control specifically at the INM, which is also termed INMAD for INM-associated protein degradation (Pantazopoulou et al., 2016). Accumulation of Mps3 leads to nuclear membrane expansion and impairs cell division (Friederichs et al., 2011; Li et al., 2017), thus demonstrating the significance of timely turnover of resident INM proteins, such as Mps3, at the nuclear envelope.

**APC/C regulates Mps3 degradation at the INM**

Four lines of evidence that we have obtained support the idea that APC/C^Cdh1^ modulates Mps3 turnover by targeting the N-terminus of Mps3. First, our combined genetic, cytological, and biochemical observations demonstrate that Cdh1, an APC/C regulator, controls Mps3 degradation. In the absence of Cdh1, Mps3 is stabilized and accumulates at the nuclear periphery. Second, inactivation of APC/C leads to increased Mps3 stability. Third, at its N-terminus, which is located in the nucleoplasm, Mps3 harbors two redundant degradation motifs that resemble the KEN and D boxes that are subject to APC/C regulation. Finally, by grafting the N-terminus of Mps3 onto an unrelated INM protein, we show that the chimeric Mps3(1-94)-Heh2 behaves in a similar manner to Mps3, responding to APC/C^Cdh1^ regulation. APC/C^Cdh1^ modulates cell-cycle progression by controlling the degradation of mitotic factors to ensure proper exit of mitosis (Visintin et al., 1997; Zachariae et al., 1998). In addition, APC/C^Cdh1^ is also active during the G1 phase in preparation for mitosis (Pines, 2011). That Mps3 is subject to APC/C^Cdh1^ regulation indicates that turnover of Mps3 is coupled to cell-cycle progression, consistent with Mps3’s mode of action in SPB duplication and separation in budding yeast (Jaspersen et al., 2002; Friederichs et al., 2011; Li et al., 2017). Our observation of Mps3 being degraded by nucleus-localized proteasomes further supports the idea that Mps3 is ubiquitinated by APC/C, which is concentrated inside the nucleus in budding yeast. To our knowledge, ours is the first example demonstrating that APC/C^Cdh1^ controls the degradation of an INM-localized protein.

How, then, does APC/C^Cdh1^ regulate the degradation of a membrane protein at the INM? We have shown that the N-terminal domain of Mps3, which is located in the nucleus, harbors two putative destruction motifs recognized by APC/C. We speculate that APC/C^Cdh1^ is recruited to the nuclear periphery to ubiquitinate Mps3, the process of which is further enhanced by protein phosphorylation at S70, as crosstalk between ubiquitination and protein phosphorylation is common for APC/C substrates (Hall et al., 2008; He et al., 2013; Davey and Morgan, 2016). Ubiquitinated Mps3 could then serve as a substrate for the Cdc48 protein complex, which extracts Mps3 from the INM and delivers it to the nuclear proteasome for degradation (Fig 8). Alternatively, we have shown previously that Mps3 is cleaved near the destruction motifs at its N-terminus (Li et al., 2017). Cleavage of Mps3 perhaps signals the critical first step in Mps3 turnover, followed by extraction of the remaining Mps3 from the INM for proteasome degradation. We note that the above two possibilities are not mutually exclusive, and future studies will clarify the process of how membrane-bound Mps3 is transported to the proteasome.

### APC/C works in concert with the E2 enzymes Ubc7 and Ubc6

Turnover of Mps3 requires the E2 conjugating enzymes Ubc7, and to some degree Ubc6, indicating that APC/C^Cdh1^ acts in concert with a set of ERAD-specific E2 ubiquitin-conjugating enzymes for proteolysis. The cognizant E2 enzymes pairing with APC/C are Ubc1 and Ubc4 (Finley et al., 2012). The nucleus-localized Ubc4 appears not to be necessary for Mps3 turnover (our unpublished data). Currently, whether Ubc1 regulates Mps3 turnover is unknown. Nevertheless, our findings have expanded the repertoire of the E3 ligases involved in ERAD-mediated membrane protein degradation, but also revealed the possibility that APC/C may pair with previously unrecognized E2 enzymes to regulate protein degradation at specialized cellular compartments, such as the INM.

Previous works have revealed that the ERAD branches formed by Doa10 and the Asi1-3 complex regulate INM protein degradation (Swanson et al., 2001; Foresti et al., 2014; Khmelinskii et al., 2014), in particular for degradation of membrane proteins mislocalized to the INM (Khmelinskii et al., 2014). However, the specificity of their degradation mechanism is unknown. In contrast, APC/C recognizes its substrates primarily through the D and KEN boxes (Davey and Morgan, 2016). Although currently is unknown whether APC/C regulates resident INM proteins other than Mps3, our findings suggest that the destruction motif(s) that reside on membrane proteins play a critical role in modulating their quality control.

### The significance of protein quality control at the INM

The SUN-domain protein Mps3 is a resident INM protein, whose anchoring at the INM is enhanced by its binding partners in the nucleus as well as the KASH-domain/like proteins bound to the ONM (Tapley and Starr, 2013). Mps3 is homologous to mammalian SUN1, whose accumulation at the INM has been implicated in progeric and dystrophic laminopathies in a mouse model (Chen et al., 2012). Similarly, accumulation of Mps3 in budding yeast leads to nuclear envelope expansion and nuclear tubule formation (Friederichs et al., 2011), and impairs cell-cycle progression (Li et al., 2017, and this study). Currently, little is known of how SUN1 is degraded in mammalian cells. Our finding that APC/C mediates Mps3 turnover during interphase of the cell cycle has implications for understanding the clearance of SUN1 and its associated proteins in animal cells, and therefore is relevant to the understanding of laminopathies in humans.

## Materials and Methods

### Yeast strains and plasmids used in this study

Yeast strains used in this study are listed in Table S1. To tag the N-terminus of Mps3 with V5, we used a homologous recombination-based gene replacement method as follows. We linearized plasmid pHG553 (see below) with AflII and integrated it at the endogenous *MPS3* locus by yeast transformation. *URA3* positive colonies were then counter-selected on a 5’-FOA plate to remove the untagged *MPS3* copy, thereby leaving *V5-MPS3* as the only functional *MPS3* gene in the yeast genome. A similar method was used to generate V5-tagged Mps3-S70A, Mps3-S70D, Mps3-3A, Mp3-2D, Mps3-3A2D, and Mps3-NC using the derivatives of plasmid pHG553. All these *MPS3* alleles were verified by DNA sequencing before use. For C-terminal tagging of Asi1 and Erg11 with 3xHA and V5, we used a standard PCR-based homologous recombination method (Longtine et al., 1998). Tub4-mApple and others have been reported previously (Shirk et al., 2011; Li et al., 2015). Gene deletion strains of *cdh1Δ, asi3Δ, doa10Δ, hrd1Δ, ubc6Δ*, and *ubc7Δ* were obtained from the ATCC deletion collection (ATCC-GSA-5). To generate the *P*_*GAL*_*-GFP-MPS3(1-94)-HEH2* fusion allele, we transformed plasmid pHG479 (see below). The following alleles, *cdc16-1, cdc20-1, cdc48-6*, and *P*_*GAL*_*-GFP-MPS3* have been reported previously (Sethi et al., 1991; Yamamoto et al., 1996; Li et al., 2017). The anchor-away strains have been described previously (Nemec et al., 2017). Briefly, rapamycin was used to force the formation of a ternary complex between an anchor protein fused with the human FK506 binding protein (FKBP12), and the target protein fused with GFP and the FKBP12-rapamycin-binding domain (FRB) (Haruki et al., 2008). In this case, the target protein was the proteasomal lid subunit RPN11 and the anchor proteins were either RPL13 or PMA1, to force localization of RPN11 to the cytoplasm or plasma membrane, respectively (Nemec et al., 2017). Due to the toxic nature of rapamycin to wild-type yeast cells, rapamycin-resistant strains containing mutated *TOR1* were used. The yeast homolog to FKBP12 is FPR1, so to reduce competition for FRB binding between FPR1 and the anchor protein, the FPR1 gene was deleted using standard PCR-based homologous recombination (Nemec et al., 2017).

Plasmids used in this study are list in Table S2. We used pRS306 as the backbone to generate *P*_*MPS3*_*-V5-MPS3* (pHG553). The *MPS3* promoter was obtained from pHG454 (Li et al., 2017). PCR-based site-directed mutagenesis was used to generate *mps3* mutant alleles (Table S2). To generate *P*_*GAL*_*-GFP-HEH2* (plasmid pHG479), the open reading frame of *HEH2* was amplified from the S288C yeast strain background and cloned into the BamHI and SacI sites of pHG302 (Li et al., 2015). To create *P*_*GAL*_*-GFP-MPS3(1-94)-HEH2* (pHG572), the first 282 bp of the *MPS3* gene was cloned into the BamHI site of pHG479. All plasmids were confirmed by DNA sequencing before use.

PCR primers used in this study are listed in Table S3.

### Yeast culture methods and yeast viability assay

For cycloheximide-chase experiments, yeast cells were grown overnight in YPD (1% yeast extract, 2% peptone, and 2% dextrose) liquid medium to saturation at 30°C. Cell cultures were diluted with YPD to reach an OD (optical density, λ = 600nm) of 0.2 and incubated at 30°C until the OD reached 0.7. Cycloheximide (CHX) was then added to a final concentration of 200 μg/mL, the point of which was defined as time zero. Cell aliquots were withdrawn at indicated times for protein extraction and prepared for western blotting.

To chemically inactivate the proteasome activity (Fig 1E), yeast cells with *pdr5Δ* were prepared as described above to reach an OD of 0.7; the culture was then split into three fractions: DMSO only, DMSO with CHX (200 μg/mL final concentration), and DMSO with CHX (200 μg/mL final concentration) and with MG132 (50 μM final concentration). Cell aliquots were withdrawn at indicated times, and protein extracts were prepared for western blotting.

To relocate the proteasomes from the nucleus to the cytoplasm or the plasma membrane (Fig 2), yeast cells containing the anchor away system (Nemec et al., 2017) were prepared as described above to reach an OD of 0.7; the culture was then split into two fractions: DMSO with CHX (200 μg/mL final concentration), and DMSO with CHX (200 μg/mL final concentration) and with rapamycin (1 μg/mL final concentration). Cell aliquots were then withdrawn at the indicated times for microscopy and protein extraction.

To arrest yeast cells at G1 (Fig 5B), strains were grown in YPD liquid medium to an OD of 0.5, then 10 μg/mL of alpha factor was added. After two hours of incubation, CHX (200 μg/mL final concentration) was added and cell aliquots collected at the indicated times for protein extraction and western blotting.

Strains carrying temperature sensitive alleles (*cdc48-6, cdc16-1, cdc20-1*) were grown overnight at 25°C to saturation in YPD liquid medium and diluted as described above with incubation at 25°C until the OD reached 0.6. The culture was then split into two fractions and allowed to incubate for 1 hour at either 25°C or 37°C. CHX was then added to a final concentration of 200 μg/mL and cell aliquots were withdrawn at the indicated times for protein extraction and subsequent western blotting (Figs 1, 5 and 7).

For experiments utilizing the *GAL1-10* promoter (Fig 7), synthetic complete (SC) medium with 2% dextrose was used to grow cells to an OD of 0.5 and an aliquot was collected for protein extraction and western blotting. Then, yeast cells were washed twice in water and incubated for about three hours in SC medium with 2% galactose to induce expression of the *GAL1-10* promoter. CHX (200 μg/mL final concentration) was added and cell aliquots collected at the indicated times for protein extraction and western blotting. To determine cell viability, yeast cells were grown overnight in YPD liquid medium to reach saturation, 10 fold diluted, spotted onto SC plates with either 2% dextrose or 2% galactose, and then incubated at 30°C for about two days (Figs 3, 5, 6 and 7).

### Genetic screen to identify *MPS3* interacting factors

From the yeast deletion collection (ATCC-GSA-5), a pooled Mata library containing deletions of individual open reading frames that encode the known E2, E3 enzymes and their regulators was used to screen *MPS3* interacting genes. This library was crossed to the yeast strain HY4671, which contains a galactose inducible *MPS3* construct, *P*_*GAL*_*-MPS3*, along with *P*_*MFA1*_*-HIS5::ade2*, which is specifically expressed in Mata cells (Tong et al., 2001). After mating, diploids were selected, grown to saturation, and incubated in sporulation medium (1% potassium acetate, 0.1% yeast extract, 0.05% dextrose) for ∼four days at 30°C. To select for haploids, sporulated cells were then transferred to SC-leucine and -histidine double dropout medium with G418 in the presence of either 2% dextrose or 2% galactose, and then incubated at 30°C for about two days to determine cell viability (Fig 3A).

### Protein extraction and western blotting

Yeast aliquots were withdrawn at the indicated times for protein extraction by precipitation in the presence of 20 mM NaOH and standard SDS-PAGE and western blotting were performed. V5-Mps3, its derivatives, and Asi1-V5 were detected by a mouse monoclonal anti-V5 antibody (1:5,000, Thermo Fisher Scientific, cat#R960-25). Erg11-3HA was detected by an anti-HA mouse monoclonal antibody (1:1,000, 12CA5, Sigma). GFP-tagged proteins were detected by an anti-GFP mouse monoclonal antibody (1:10,000 dilution, Thermo Fisher Scientific, cat#GF28R). The level of Pgk1 was probed by a Pgk1 antibody (Thermo Fisher Scientific, cat#PA5-28612) to serve as a loading control. Horseradish peroxidase-conjugated secondary antibodies, goat anti-mouse (Bio-Rad, cat#1706516), were used to probe the proteins of interest by an enhanced chemiluminescence (ECL) kit (Bio-Rad, cat#1705060).

Two ECL-based western blot detection methods were utilized, X-ray film (Figures 1-7 and S1) and the ChemiDoc MP Imaging System (Bio-Rad, cat#17001402) (Figure S2). To calculate the relative protein abundance at each time point, individual band intensities were measured using the IPLab Imaging Software in conjunction with an in-house GelAnalyzer script and exported to Microsoft Excel. Target protein band intensities were made relative to those of the loading control (Pgk1).

### Fluorescence Microscopy

A DeltaVision microscope was used to acquire live-cell fluorescence images as described previously (Li et al., 2015). Briefly, an agarose pad with 2% raffinose was prepared on a concave slide and an aliquot of yeast cells was placed on the agarose. The slide was then sealed with LVP and images were acquired at each of the indicated time points. Images were acquired with a 63x (NA=1.40) objective lens on an inverted microscope (IX-71, Olympus) equipped with a CoolSNAP HQ2 CCD camera (Photometrics). For images attained during time course experiments, 10 optical sections with a 0.5 μm thickness were acquired at each time point. To reduce phototoxicity, a neutral density filter was used to diminish the intensity of the excitation light to about 50% or less of the equipment output.

## Acknowledgements

We thank Dr. Yanchang Wang for insightful discussion and for providing plasmids and yeast strains. Elizabeth Staley provided technical assistance; Jen Kennedy assisted with text editing. This work is supported by the National Institute of General Medicine of the National Institutes of Health under award number GM117102.

---

**Supplemental Figure 1.**
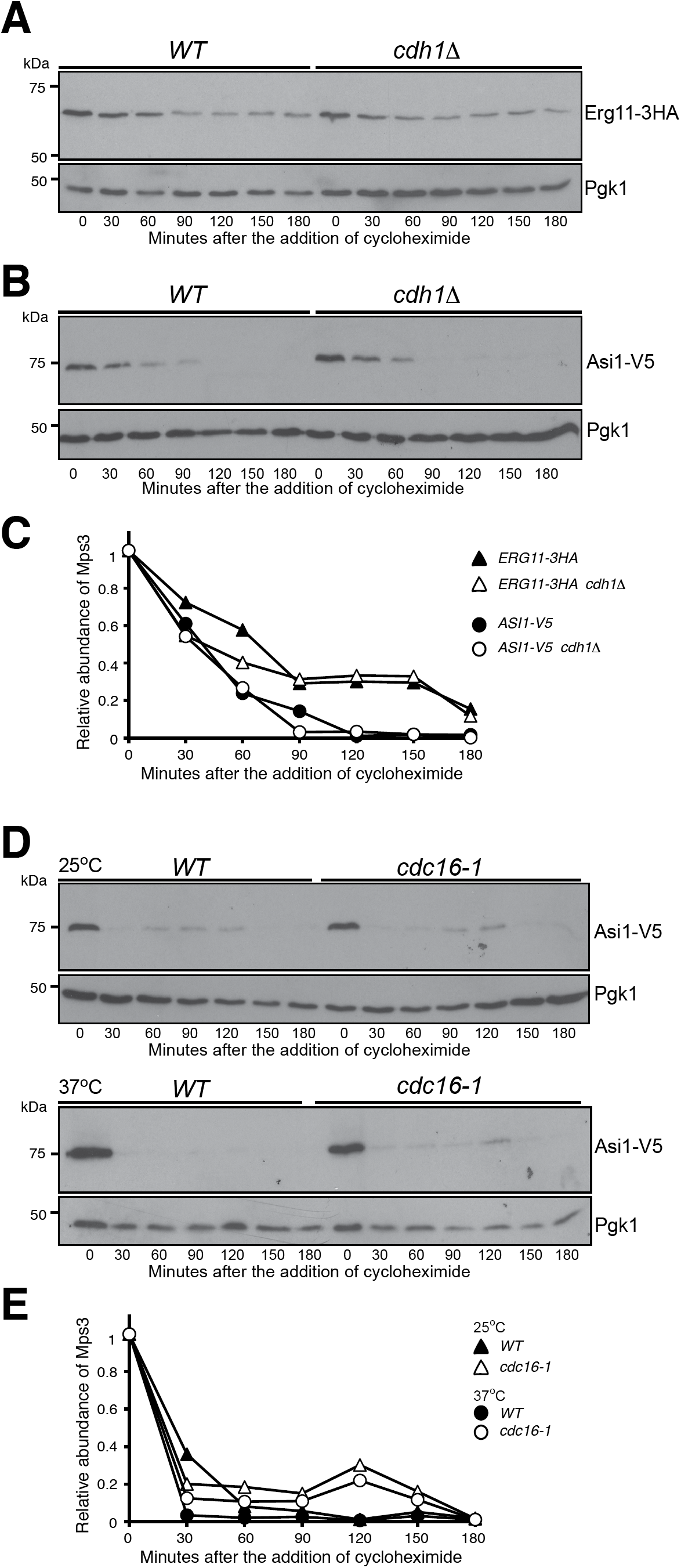
Degradation of Erg11 and Asi1 is independent of Cdh1 and APC/C. **(A-C)** Degradation of Erg11 and Asi1 is independent of Cdh1. The relative protein abundance of Erg11 (**A**) and Asi1 (**B**) in *WT* and *cdh1δ* cells was analyzed after arresting cells in G1 using alpha factor (10 μg/mL) for 2 hours and then performing CHX chase as described in Figure 5B. Quantification of the relative protein abundance is shown in panel C. **(D** and **E)** Degradation of Asi1 is independent on Cdc16. Cells were grown at 25°C to the exponential phase and CHX was then added as shown in Figure 5E. To inactivate *cdc16-1*, cells were shifted to 37°C for 1-hour before the addition of CHX. Quantification of Asi1-V5 protein abundance is shown in panel E. Note there is no change in stability between the permissive and nonpermissive temperatures.

**Supplemental Figure 2.**
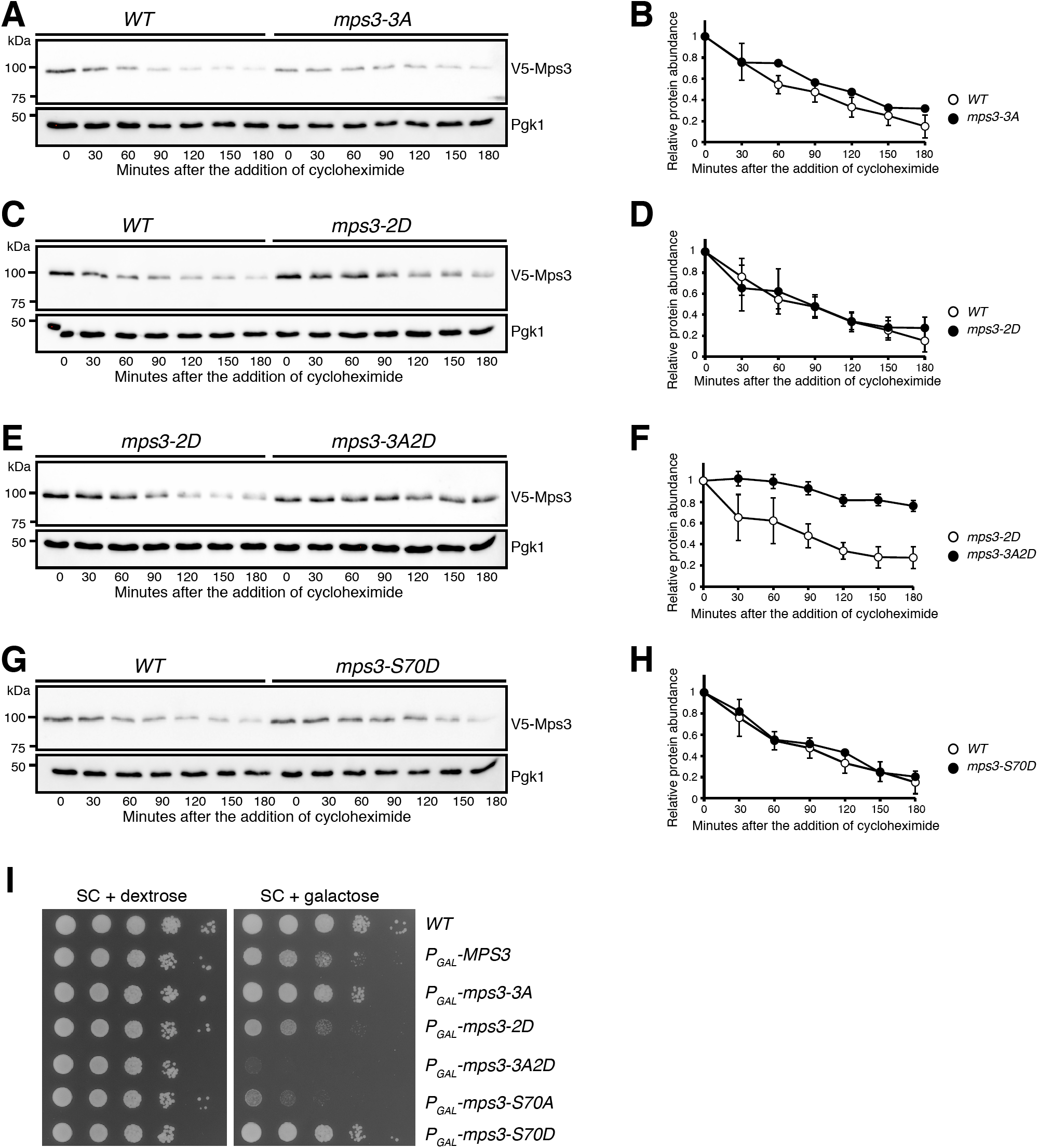
Redundant destruction motifs located at the N-terminus of Mps3. **(A** and **B)** The effect of the KEN box on Mps3 protein stability. Yeast cells were prepared for CHX chase and analyzed as described in Figure 1A. Quantification of Mps3 protein abundance is shown in panel B. (**C** and **D**) The effect of the putative D box on Mps3 protein stability. Quantification of Mps3 protein abundance is shown in D. (**E** and **F**) The KEN and D boxes are redundant in regulating Mps3 degradation. Quantification of Mps3 protein abundance is shown in F. (**G** and **H**) Phosphorylation at S70 plays a role in Mps3 stability. Quantification of Mps3 protein abundance is shown in H. **(I)** Overproduction of stabilized Mps3 causes cell lethality. Yeast cells were grown overnight in YPD liquid medium to reach saturation, 10 fold diluted, spotted onto SC plates with either 2% dextrose or 2% galactose, and then incubated at 30°C for about two days.

**Supplemental Figure 3.**
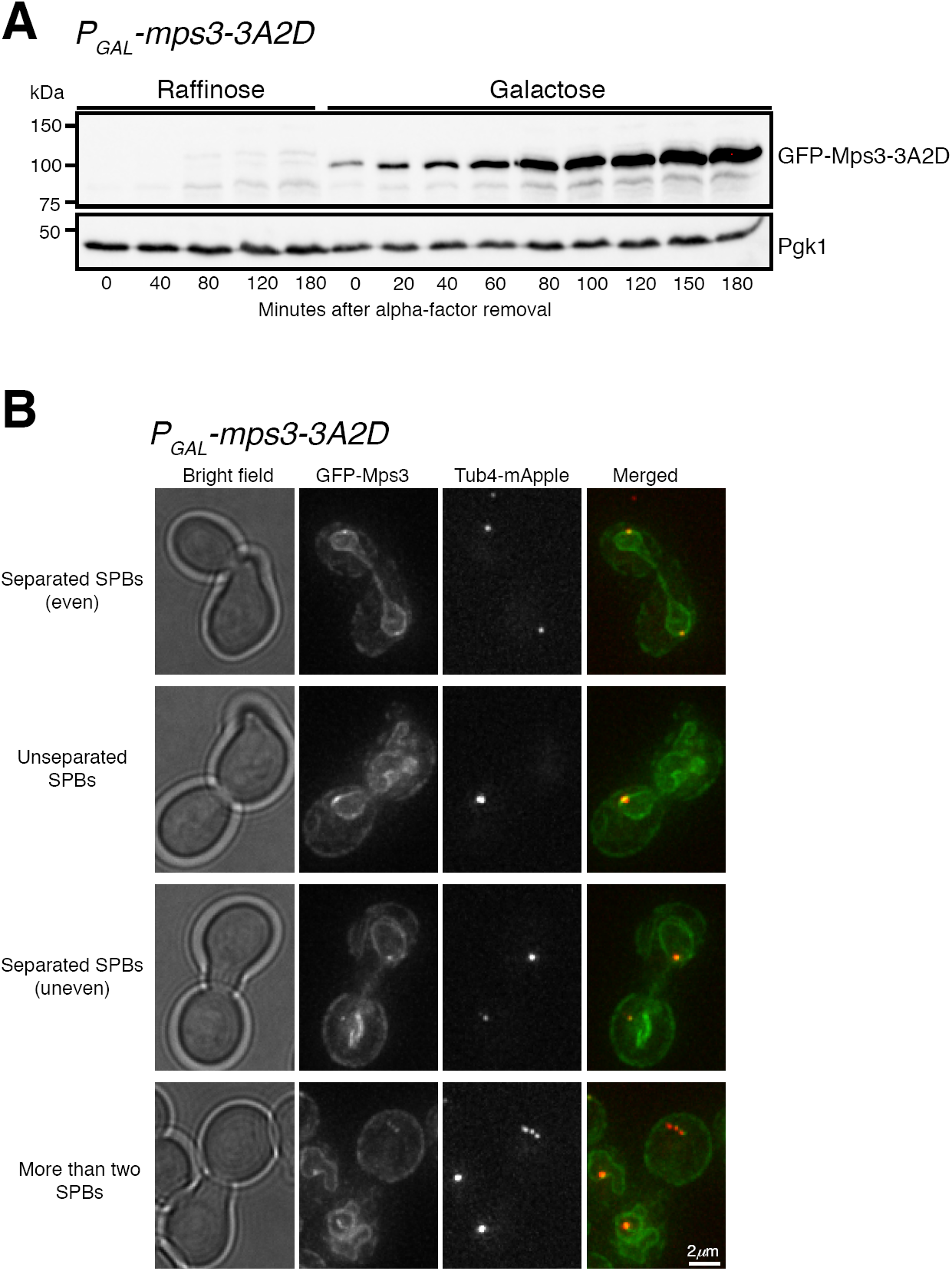
Overexpression of *mps3-3A2D* leads to defective SPB separation. **(A)** Protein level of Mps3-3A2D upon galactose induction of *P*_*GAL*_*1-mps3-3A2D*. Yeast cells were grown in raffinose and arrested at G1 with alpha-factor as shown in Figure 6I. To induce the *GAL1-10* promoter, galactose was added to the culture medium 30 minutes prior to the removal of alpha-factor. Cell aliquots were withdrawn at the indicated times and prepared for western blot. Time zero refers to the point of alpha-factor removal. The level of Pgk1 serves as a loading control. **(B)** Representative images showing the localization of GFP-Mps3 and Tub4-mApple. Aliquots were withdrawn and prepared for live-cell fluorescence microscopy. Tub4-mApple marks the SPB. Four categories of SPB separation in large budded cells were classified, the first being normal SPB separation in which SPBs separated evenly, the remaining three being types of defective SPB separation: unseparated SPBs (one Tub4-mApple spot), uneven separation of SPBs (two unequal Tub4-mApple spots), and more than two SPBs (more than 2 Tub4-mApple spots in a single cell).

Supplemental Table S1. Yeast strains used in this study.

Supplemental Table S2. Plasmids used in this study.

Supplemental Table S3. Primers used in this study.

